# MetaUmbra: Statistically Controlled Genome-Level Presence Inference from Metaproteomic Peptides

**DOI:** 10.64898/2026.04.29.721689

**Authors:** Qing Wu, Zhibin Ning, Ailing Zhang, Kai Cheng, Daniel Figeys

## Abstract

Taxonomic interpretation of metaproteomic peptides remains difficult because many peptide sequences are present in proteins from different organisms, reducing taxonomic specificity. Current peptide-centric workflows can report taxonomic summaries or taxon-level confidence scores, but they do not directly provide formal genome-level statistical inference for user-defined genome collections. Here we present MetaUmbra, a tool that derives genome-level statistical evidence from identified peptides. MetaUmbra builds theoretical peptide lists by *in silico* digestion of genome proteins, matches observed peptides against these references, and evaluates unique and shared peptide evidence within a formal statistical framework that accounts for peptide sharing across the reference panel. In the defined community benchmark SIHUMIx, MetaUmbra recovered the expected genomes without additional significant calls after embedding them in a large gut reference background. In the single-strain benchmark Mix24X, all expected genomes were recovered under near-neighbor and full-background expansion, with additional calls mainly associated with closely related genomes. In a hamster gut genome panel, MetaUmbra further preserved an interpretable ranking of candidate genomes in a dense real-data setting. Together, these results show that MetaUmbra provides statistically controlled genome-level presence inference while accounting for ambiguity arising from shared peptides. MetaUmbra therefore provides a practical framework for converting peptide evidence into genome-level statistical inference in metaproteomics.

## Introduction

Metaproteomics measures functional changes at a community level by identifying and quantifying peptides using mass spectrometry. It is now widely applied across diverse environments, including the human gut, soil, ocean, and other host-associated microbiomes (1, 2). A persistent challenge in metaproteomics is to provide statistical evidence of the presence of specific genomes or strains using identified peptides, because many peptides originate from conserved proteins and therefore map to multiple related organisms. This challenge becomes more pronounced as reference collections expand and as analyses move from generic databases to larger custom panels (3, 4).

Current workflows approach this problem in different ways. Protein-centric approaches first infer proteins and then annotate taxa, whereas peptide-centric approaches work directly from peptide evidence (1, 5). For example, Unipept (6) applies the lowest common ancestor rule to peptide taxonomic assignment. This is robust because it avoids overly specific assignments when peptides are shared, but it also reduces resolution by moving ambiguous evidence to higher taxonomic ranks and does not use peptide scores in the inference step. Protein-centric workflows such as MetaLab (5) can gain specificity through protein inference, but they introduce additional assumptions and potential bias from protein grouping. Such outputs are useful for taxonomic overview, but they are not designed to test whether a specific genome in a user defined panel is present with formal genome-level statistical support.

Shared-peptide ambiguity is also a long-standing problem in protein inference. ProteinProphet (7) and related probabilistic methods (8, 9) use peptide-to-protein relationships to evaluate candidate proteins, while Pipasic (10) accounts for similarity among reference proteomes in strain-level abundance estimation. These approaches address related ambiguity problems, but their protein-inference or abundance objectives differ from genome-level presence testing within a user-defined reference collection.

Some methods add quantitative confidence to peptide derived taxonomic interpretation. Peptonizer2000 (11) uses Bayesian inference to generate taxon level confidence scores from peptide lists and can operate at user selected taxonomic levels. MiCId (12) reports species or genus level statistical confidence, but it is a self contained workflow that couples peptide identification with taxonomic inference through its own database search strategy. These methods are useful, yet they do not provide genome-level p-values and q-values from peptide lists for user defined genome collections. This gap is important for applications based on custom strain panels, metagenome assembled genome catalogs, combined host microbe references, or iteratively refined follow up panels.

Reference-database composition influences both peptide identification and downstream taxonomic interpretation in metaproteomics (3, 13). Recent DIA metaproteomics studies have demonstrated deep and reproducible profiling of complex microbial communities (14, 15), while sequencing-based microbiome workflows such as QIIME 2 provide complementary taxonomic and community-level context (16). These advances address important aspects of reference selection, data acquisition, and community profiling, but they do not directly resolve genome-level presence inference from peptide evidence shared across closely related organisms.

Here we present MetaUmbra, a peptide-centric workflow for genome-level presence inference from metaproteomic peptide lists. MetaUmbra builds theoretical peptide lists by *in silico* digestion of genome proteins, matches observed peptides to these references, and evaluates unique and shared peptide evidence within a formal statistical framework. Shared peptide evidence is assessed against an empirical null model by Monte Carlo sampling and then combined with unique-peptide evidence to obtain genome-level significance values. We evaluated the method using two complementary benchmarks. SIHUMIx (4) is a defined mixed community composed of eight expected gut genomes and was used to test recovery in a controlled community background. Mix24X (17) comprises 24 individually measured strains and was used to test genome discrimination under stronger peptide sharing and near-neighbor interference. Together, these benchmarks assess recovery in mixed communities and discrimination among individual strains under expanded reference backgrounds. In this study, we used a representative human gut genome background derived from the UHGG (18) catalogue of MGnify (19) for benchmarking. The same framework can be applied to other user-defined genome collections, although the resulting inference should be interpreted relative to the composition of the reference collection.

## Experimental Procedures

### 1. MetaUmbra workflow overview

MetaUmbra performs genome-level presence inference from identified peptide lists against a user-defined genome collection through two main components: reusable reference construction and genome presence scoring. Protein FASTA files from the genome collection are first digested *in silico* to generate genome-specific theoretical peptide reference tables. This reference construction step only needs to be performed once for a given genome collection and digestion setting. Observed peptides are then matched against these references and evaluated within a formal statistical framework. Shared peptide evidence is weighted by peptide degeneracy across the tested genome panel and assessed against an empirical null distribution by Monte Carlo sampling. Unique peptide evidence is evaluated separately, and the two evidence components are then combined to obtain a genome-level p-value. The resulting genome-level p-values are adjusted across candidate genomes using the Benjamini–Hochberg procedure (20).

### 2. Construction of genome-specific theoretical peptide reference databases

For each user-defined genome collection, protein sequences were organized as genome-specific FASTA files, with one FASTA file representing one genome. Each genome FASTA was subjected to *in silico* digestion using Rapid Peptides Generator (RPG, v2.0.5) (21) under digestion settings chosen to match the upstream peptide identification workflow. For the benchmark analyses reported here, digestion used Trypsin/P, peptide lengths of 7–30 amino acids, and up to two missed cleavages. Retained peptides were exported for each genome as peptide reference tables linking peptide sequences to their source proteins. These genome-specific peptide tables formed the theoretical reference database used by MetaUmbra. Representing the reference in this genome-resolved manner preserves the link between peptides and candidate genomes, supports direct matching of observed peptides, and provides the basis for quantifying peptide degeneracy across the tested genome collection.

Because conventional tandem mass spectrometry generally does not distinguish isoleucine from leucine, the primary analyses used I/L-equivalent peptide matching. I and L were converted to a common representation in both observed and theoretical peptides before deduplication, matching, and degeneracy calculation. Exact-sequence matching was retained as a sensitivity analysis for the hamster dataset; Q/K and AD/W were not collapsed. Reference collections should include plausible close competitors when genome-level discrimination is required. For complex microbiomes, matched metagenome-derived genomes or MAGs are preferred when available, whereas ecologically relevant public genome catalogues can be used when matched sequencing data are unavailable.

### 3. Genome presence scoring from an observed peptide list

Observed peptides were supplied as identified peptide tables from workflows such as MaxQuant (22) or DIA-NN (23). MetaUmbra operates at the peptide-sequence level. Available decoy and peptide-error fields were used to filter identifications before scoring. I/L normalization was applied before deduplication, and duplicate normalized sequences were collapsed by retaining the highest available evidence score. For each scoring run, retained sequence scores were min-max scaled to [0,1] as

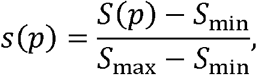

where *s*_min_ and *s*_max_ are the minimum and maximum retained scores. If no informative score was available, retained peptides were assigned *s*(*p*) - 1. These scores were used as relative evidence weights rather than identification probabilities. Peptide-error measures were kept separate from score weights and from the fixed α = 0.05 used by the unique-evidence models.

For genome *k*, let *τ*_k_ denote its theoretical peptide set and *?* the observed peptide set after preprocessing. The matched peptide set was defined as

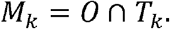

To account for peptide sharing across the tested genome panel, each observed peptide *p* was assigned a degeneracy *d*(*p*), defined as the number of genomes in the panel containing *p*. Each matched peptide was weighted as

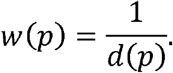

Thus, peptides unique to one genome retained full weight, whereas shared peptides were down-weighted according to their degeneracy. For each genome, weighted evidence was summarized as

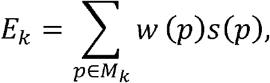

where *s*(*p*) denotes the retained relative peptide evidence score. Shared peptide evidence was defined by matched peptides with *d*(*p*) > 1, whereas unique peptides had *d*(*p*) - 1.

Shared peptide significance was evaluated using a peptide-space Monte Carlo empirical null. Weighted contributions from observed shared peptides, *w*(*p*)*s*(*p*), were pooled by peptide degeneracy bin (2-5, 6-20, 21-100, 101-500, and > 500). These bins represent broad peptide-sharing regimes and were not optimized against benchmark outcomes. For each genome, a genome-specific null distribution was generated by sampling from these pools according to that genome’s matched shared-peptide bin counts. The empirical shared-evidence p-value was calculated as

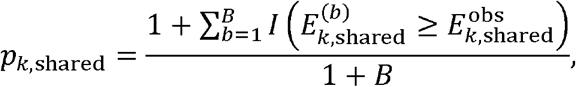

where *I* (·) is the indicator function, 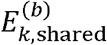 is the shared evidence obtained in Monte Carlo replicate *b*, 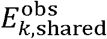 is the observed shared evidence, and *B* is the number of replicates. This procedure evaluates whether the observed shared support exceeds that expected from ambiguity-aware peptide reuse without requiring a separate decoy-genome database. Component scoring used 500 first-stage replicates with seed 1. Near-threshold refinement used 20,000 replicates for the individual Mix24X, pooled near-genus, and hamster analyses, and 200,000 replicates for SIHUMIx and pooled full-UHGG Mix24X.

Unique evidence accounts for differences in theoretical panel-unique peptide opportunity. For genome *k,A*_*k*_ denotes the number of distinct theoretical peptides assigned only to that genome within the active reference panel, calculated after peptide normalization and before observed-peptide scoring. In the empirical-background model, the observed unique-peptide count *?*_*k*_ was compared with weak-background genomes having similar *A*_*k*_. The candidate genome itself and the highest-support candidates were excluded from this background. If *Q*_0.95,*k*_ denotes the 95th percentile of unique-peptide counts in the comparable background, the unique-evidence statistic was

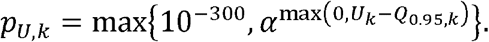

The numerical floor was 10^−300^, and α = 0.05. When an adequate comparable background was unavailable, the alpha-upper-bound model instead used

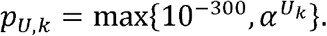

This multiplicative model treats α as an upper bound on retained unique-peptide error contributions; its interpretation therefore depends on peptide-dependence and error-bound assumptions and should not be interpreted as a posterior probability of genome presence. Automatic model selection used theoretical-opportunity strata, requiring at least 100 comparable genomes, an expected upper-tail count of at least 5, and adequate background for at least 90% of candidates. The large SIHUMIx and full-UHGG Mix24X panels used the empirical-background model, whereas the 24-genome individual Mix24X and 73-genome near-genus panels used the alpha-upper-bound model. Hamster analyses explicitly used the alpha-upper-bound model.

For the empirical-background model, the fraction of high-support genomes excluded from background estimation was updated iteratively from the observed evidence. The reported analyses used an initial exclusion fraction of 0.10. At each iteration, genomes with at least one observed peptide match were ranked by unique-peptide count, with ties resolved by unique weighted evidence, total weighted evidence, and genome identifier. The highest-ranked fraction was excluded, unique statistics were recalculated, and intermediate genome-level p-values were combined and BH-adjusted. The exclusion fraction for the next iteration was defined as the proportion of matched genomes with intermediate *q* ≤ 0.20, bounded to 0.05,0.20J. Updating stopped when the absolute change in the exclusion fraction was < 0.01 or after five iterations. The intermediate q-value threshold was used only for background estimation and did not define formal genome-presence calls. Subsequent Monte Carlo refinement updated the shared-evidence and final genome-level p-values without repeating background selection. Additional rounding, binning, and fallback rules are described in the Supplementary Methods.

The unique- and shared-evidence p-values represent complementary aspects of peptide support and were therefore combined for genome-level inference. Because both components are derived from the same observed peptide set and reference panel, independence between them cannot be guaranteed. Fisher’s method combines multiple component p-values into a single significance measure, with its standard chi-squared calibration assuming independent component p-values (24), and was therefore retained only as a sensitivity analysis. For the primary analysis, the unique- and shared-evidence component p-values were combined using the procedure defined below.

The unique- and shared-evidence component p-values, *p*_*U*_ and *p*_*S*_, were first evaluated jointly using a two-component Simes intersection test (25). The final genome-level p-value was then obtained by closed testing of the unique-evidence component (26):

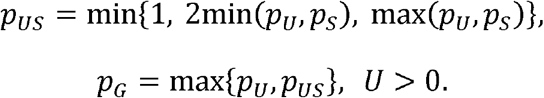

For genomes without an observed unique peptide (**U=**0), the final genome-level p-value was set to *p*_*G*_ - 1. Thus, *p*_*G*_ cannot be smaller than *p*_*U*_, and shared evidence alone cannot establish a formal genome-presence call. Genome-level p-values were adjusted using the Benjamini-Hochberg procedure (20), separately within each independently analyzed peptide set and candidate panel and across genomes with at least one observed peptide match. The primary statistical outputs are raw p-values and BH-adjusted q-values, and the presence score used for visualization is

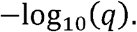

Harmonic-mean, Bonferroni, and unique-only alternatives were also retained for sensitivity comparison.

### 4. Benchmark datasets and comparative tools

MetaUmbra was evaluated using the SIHUMIx (4) and Mix24X (17) benchmark datasets because they test complementary aspects of genome-level inference from metaproteomic peptides. SIHUMIx is a defined mixed-community benchmark composed of eight expected gut genomes and was used to assess recovery in a controlled multispecies background. Mix24X is a complementary 24-species benchmark containing both phylogenetically distant species and difficult near-neighbor cases, making it suitable for testing genome discrimination and ranking stability under stronger peptide sharing. The 4,744 representative genomes from the UHGG v2.0.2 catalogue of MGnify (19) were used as the representative human gut background.

For SIHUMIx, the eight expected genomes were embedded in a species-deduplicated UHGG background. UHGG representative genomes belonging to the same species as any SIHUMIx member were removed before panel construction to avoid direct species-level redundancy between benchmark targets and background references.

For Mix24X, three complementary settings were analyzed. First, each of the 24 individual-strain peptide sets was scored against the exact 24 Mix24X genomes. Second, a synthetic pooled peptide set generated by merging the peptide sets from all 24 strains was scored against the 24 expected genomes together with a near-genus competitor panel derived from the UHGG representative set. Third, the same pooled peptide set was scored against the 24 expected genomes together with a species-deduplicated full UHGG representative background. Species-level duplicates between Mix24X targets and UHGG representatives were removed before constructing the expanded panels.

Performance was assessed by recovery of expected genomes, rank position, genome-level p-values, BH-adjusted q-values, and additional significant genomes. The primary significance threshold was *q* ≤ 0.01, with *q* ≤ 0.05 used as a secondary threshold. Additional calls were defined relative to the declared benchmark membership and were reported separately from estimated q-values. Expected benchmark members are listed in Supplementary Table S1.

For descriptive comparison, previously generated Unipept 6.4.3 (6) and Peptonizer2000 (11) outputs for the same SIHUMIx and Mix24X peptide datasets were retained. These external tools were not rerun after revision of the MetaUmbra scoring procedure, and the comparison was therefore used to provide context for expected-member recovery and output compactness rather than as an equivalent statistical benchmark.

Primary benchmark peptide datasets were obtained from the PRIDE repository (27) under identifiers PXD023217 for SIHUMIx and PXD005728 for Mix24X. SIHUMIx data were acquired by DDA LC-MS/MS and processed using MaxQuant 2.6.7.0/Andromeda, whereas Mix24X data were acquired by DDA and searched using Mascot Daemon 2.5.1. Acquisition methods, peptide-input fields, analysis units, and candidate-panel sizes are summarized in Table 1.

**Table 1.** sAcquisition, peptide-input fields, and analysis units for the primary datasets.

| Dataset | Acquisition/search | Score field | Analysis unit | Candidate panel |
| --- | --- | --- | --- | --- |
| SIHUMIx | DDA; MaxQuant 2.6.7.0 | Score (Andromeda); min-max scaled | Defined-community peptide set | 4,746 genomes |
| Mix24X | DDA; Mascot Daemon 2.5.1 | Score (Mascot); min-max scaled | 24 individual sets or one synthetic pooled set | 24, 73, or 4,764 genomes |
| Hamster | DIA-PASEF; DIA-NN 1.9.1 | score (from Evidence); min-max scaled | 58 runs pooled as one study-level union | 706 MAGs |

To assess whether MetaUmbra performance was preserved beyond the primary benchmarks, two additional published mixture datasets were included as supplementary evaluations. The first was a six-strain culture-mixture dataset from a metagenome-informed metaproteomics study (PXD057701), in which six bacterial cultures were normalized by optical density and combined into isovolumic mixtures; the original study also included 14 absent gut bacterial species in the search database to evaluate erroneous identifications (28). The second was the ATCC 20 Strain Even Mix Whole Cell Material microbiome standard analyzed in the SPEED study (PXD011189) (29). For both datasets, the source-study reference genomes were analyzed together with the UHGG representative background using the same background-expansion strategy applied to the primary benchmarks.

### 5. Application to a hamster gut reconstructed genome panel

To evaluate MetaUmbra in a real-data setting beyond the defined benchmarks, we analyzed a published hamster gut microbiome resource generated in companion SARS-CoV-2 infection studies (30). Ninety shotgun metagenomes were combined by co-assembly and binning, yielding 926 bins.

Among these, 706 bacterial bins annotated by GTDB-Tk (31) were retained as candidate genomes. These reconstructed genomes are referred to below as metagenome-assembled genomes (MAGs). Protein FASTA files were organized at the bin level and converted into genome-specific theoretical peptide reference tables using the digestion framework described above, retaining tryptic peptides of 7–30 amino acids. The published upstream MetaX search database also included downloaded complete genomes; these were not included in the 706-MAG MetaUmbra candidate panel.

Metagenome-derived reference files for this application are available at Zenodo (32) (record 13909289), and the matched hamster fecal metaproteomics dataset is available through the PRIDE repository under identifier PXD057158.

Observed peptides were derived from the corresponding hamster fecal DIA-PASEF dataset processed with DIA-NN 1.9.1. The retained report contained 58 runs whose rows already satisfied Q.Value ≤0.05; no retained decoy flag was available. Duplicate sequences were collapsed by retaining the highest Evidence score, which was used as a relative support measure rather than as an identification probability. The reduced scoring input contained Sequence, score (from Evidence), and Q.Value, and scores were min–max scaled as described in Section 3. Pooling all runs into a study-level peptide union yielded 132,106 exact sequences and 130,835 sequences after I/L normalization. The I/L-equivalent union was used for the primary analysis, with exact-sequence matching retained as a sensitivity analysis. For contextual comparison, mean transcripts per million (TPM) was calculated for each genome across the matched hamster metagenomic samples. TPM was used only as an external abundance descriptor and was not included in genome selection, statistical inference, or figure ordering.

### 6. Experimental Design and Statistical Rationale

The evaluation strategy was designed to test MetaUmbra in settings that reflect its intended use for genome-level presence inference from metaproteomic peptide lists. SIHUMIx and Mix24X provide defined expected members and enable assessment of expected-genome recovery, ranking, and additional significant calls under controlled reference expansion. The hamster MAG panel was included to assess evidence structure in a complex reference space without a closed expected set; its pooled analysis represents study-level peptide support rather than per-sample prevalence. Raw genome-level p-values and BH-adjusted q-values were reported separately from observed benchmark outcomes. The primary threshold was q ≤0.01, with q ≤0.05 used for secondary review.

Sensitivity analyses assessed the stability of formal calls to Monte Carlo settings and reference composition. We reran 27 combinations of three benchmark panels, three degeneracy-bin schemes, and random seeds 1, 7, and 42, together with eight SIHUMIx and 24 Mix24X near-genus leave-one-reference-out scenarios, using MetaUmbra 1.5.0 and with all other analysis settings consistent with the corresponding primary analyses. Default, coarse, and fine bin edges were [1, 5, 20, 100, 500], [1, 20, 100, 500], and [1, 2, 5, 10, 20, 50, 100, 500], respectively. Leave-out analyses used the default bins and seed 1, with Monte Carlo iteration counts as specified in Section 3. Identified peptide inputs were held fixed, and reference removal was performed before recalculating degeneracy, unique/shared classification, theoretical opportunity, and scoring. These analyses therefore assess scoring-stage reference incompleteness rather than an upstream database search performed with an incomplete reference.

For leave-out analyses, the baseline was the configured panel before reference omission. Peptide counts represented distinct sequences after I/L-equivalent normalization within the corresponding pooled analysis unit. Additional calls were genomes outside the expected genome-ID set in Supplementary Table S1. A newly significant call was defined as a candidate with a baseline q-value above the stated threshold and an omission q-value at or below that threshold. One call event represents one genome in one omission scenario; repeated events are therefore not independent biological observations. Matched, panel-unique, and shared peptide counts and component p-values were compared with the same genome in the baseline. Paired records are provided in Supplementary Table S5.

To verify the sequence-level basis of reference sensitivity, a peptide-to-genome mapping was constructed from the retained matched-peptide caches, and panel-unique sequence sets were enumerated before and after each omission. For sequences that gained uniqueness, the complete baseline owner set was examined. Reconstructed matched, unique, and shared counts were reconciled with the fresh baseline and all leave-out outputs. Sequence-level records are provided in Supplementary Table S6.

Significant candidates were also summarized by observed unique-peptide count (0, 1, 2, 3, or ≥4) at q≤ 0.01 and q ≤ 0.05, separating expected and additional calls in the defined benchmarks. The 24 individual Mix24X analyses were summarized as candidate-within-analysis records. Hamster I/L-equivalent and exact-sequence analyses were reported separately because their candidate sets overlap and neither has a closed truth set. The 1–3 unique-peptide category was used descriptively and was not treated as an additional calling threshold (Supplementary Table S7).

Reference-preparation cost was evaluated separately from genome scoring. We randomly sampled 500 protein FASTA files from an available 4,516-genome UHGP collection (seed 42) and selected nested 100-, 200-, and 500-genome subsets. Each subset was digested once with 16 worker processes using MetaUmbra 1.5.0, RPG 2.0.5, RPG Trypsin enzyme ID 42, up to two missed cleavages, and peptide lengths of 7–30 residues. Wall time included process startup, digestion, and TSV writing; peak summed process-tree resident memory was sampled every 0.1 s. Network transfer, input staging, output validation, peptide identification, and genome scoring were excluded. These measurements characterize implementation cost for the tested machine and workload rather than reconstruction of the historical reference digestion. Detailed settings and measurements are provided in Supplementary Table S8.

## Results

### 1. Workflow overview of MetaUmbra

Figure 1 summarizes the MetaUmbra workflow and provides the context for the benchmark results shown below. Protein FASTA files are first converted into genome-specific theoretical peptide references, after which observed peptides are matched and evaluated within a formal statistical framework to produce genome-level p-values, BH-adjusted q-values, and presence scores. To support routine use, MetaUmbra was implemented as a standalone tool with graphical user interface, command-line interface, and function-based workflows. An overview of the graphical user interface is provided in Supplementary Figure S1.

**Figure 1.**
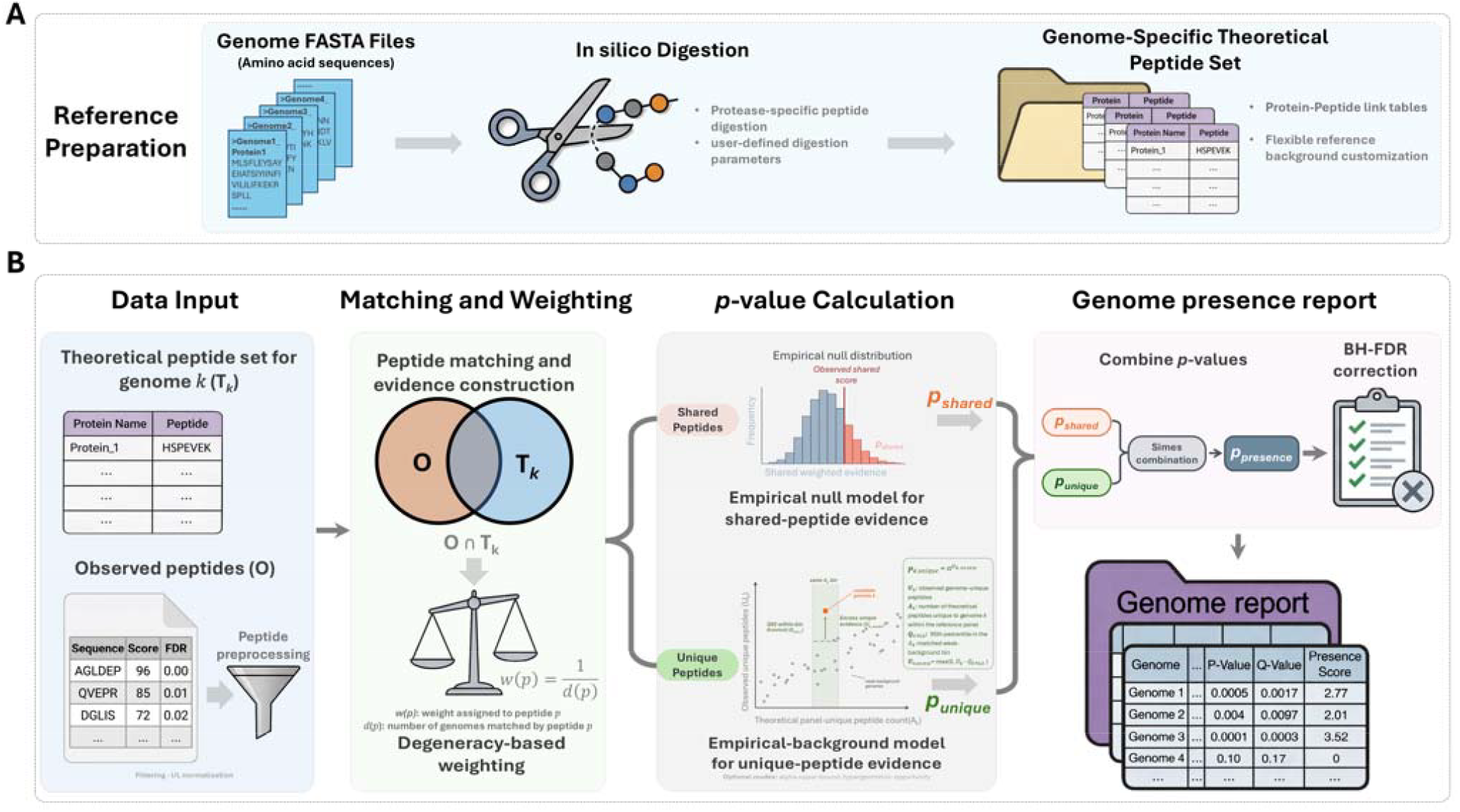
Workflow overview of MetaUmbra. (A) Protein FASTA files are digested *in silico* with Rapid Peptides Generator to build genome-specific theoretical peptide references. (B) Observed peptides are matched against these references and evaluated within a formal statistical framework to produce per-genome p-values, BH-adjusted q-values, and presence scores.

### 2. Theoretical peptide reference space defines the ambiguity landscape

To characterize the ambiguity that MetaUmbra must resolve, we performed *in silico* digestion of the 4,744 representative genomes in the UHGG collection using the digestion settings applied in the benchmark analyses. This analysis quantifies peptide sharing in the reference space itself, before any observed peptides are considered, and therefore defines the ambiguity landscape for downstream inference. Figure 2 summarizes this sequence-level theoretical reference space; subsequent benchmark analyses use I/L-equivalent peptide matching as described in the Experimental Procedures.

**Figure 2.**
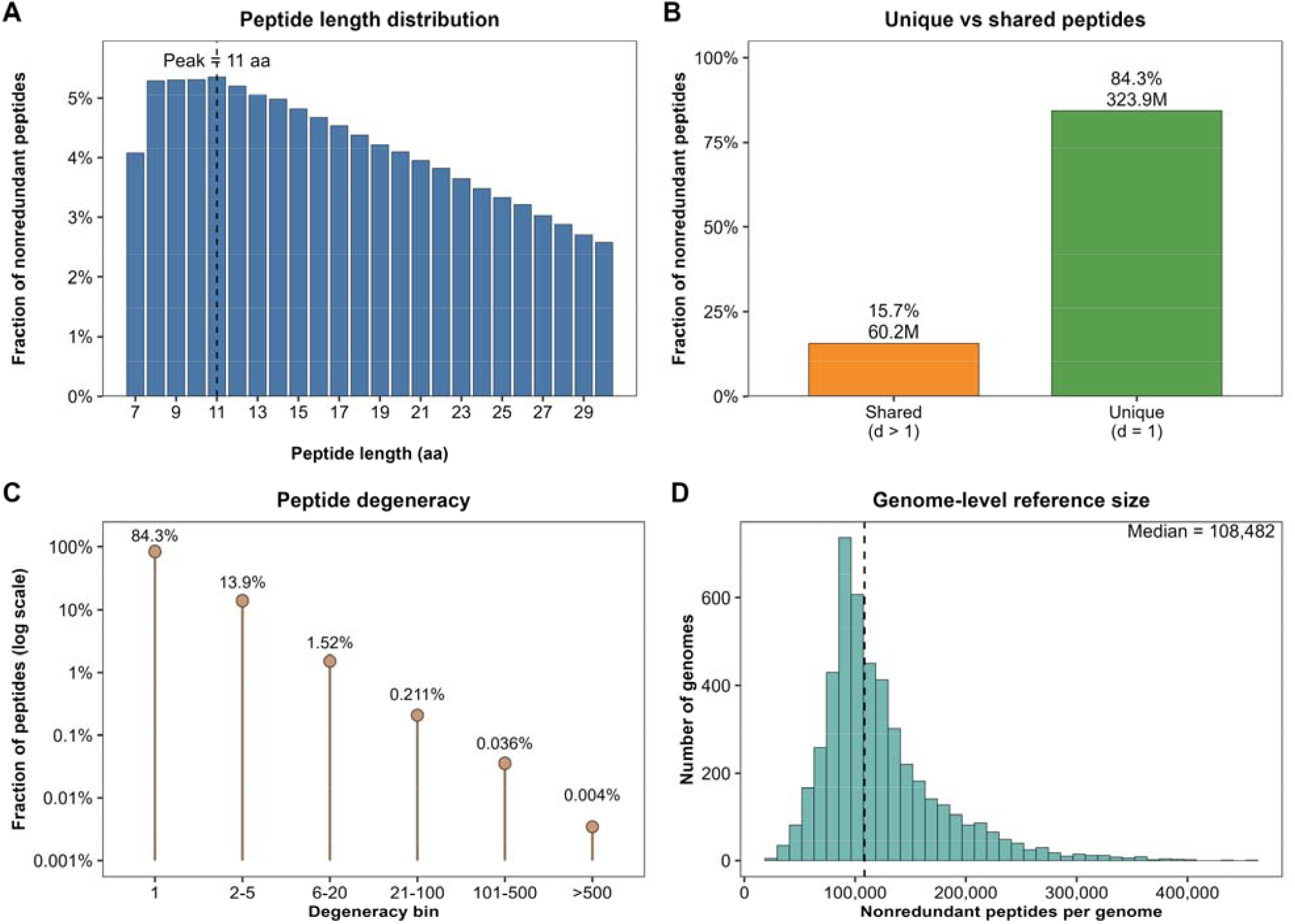
Characterization of the UHGG theoretical peptide reference space. (A) Length distribution of nonredundant theoretical peptides generated by *in silico* digestion of 4,744 genomes. (B) Fractions of unique and shared nonredundant theoretical peptides. (C) Distribution of reference-level peptide degeneracy, showing that most sharing occurs at low degeneracy orders. (D) Distribution of nonredundant theoretical peptide counts per genome.

The resulting theoretical peptides showed the expected length distribution under the selected digestion settings (Figure 2A). Across the 4,744 genomes, the digest produced 384.1 million nonredundant theoretical peptide sequences. Among these, 323.9 million (84.3%) were unique to a single genome, whereas 60.2 million (15.7%) were shared across at least two genomes (Figure 2B). Thus, although genome specific peptides predominated, peptide sharing remained substantial at the scale of the UHGG background and could not be ignored during downstream inference.

To examine the structure of peptide sharing, we summarized theoretical peptides by peptide degeneracy, defined as the number of genomes containing a given peptide. Most shared peptides were concentrated in low order bins (Figure 2C). Specifically, 13.9% of all nonredundant peptides fell into the 2–5 bin and 1.52% into the 6–20 bin, whereas only 0.21% exceeded 20 genomes, 0.036% fell into the 101–500 bin, and 0.004% were shared by more than 500 genomes. Peptide ambiguity in the UHGG-derived reference space was therefore substantial but predominantly local rather than global.

We also observed substantial heterogeneity in genome specific reference size. The number of nonredundant theoretical peptides per genome showed a broad right skewed distribution, with a median of 108,482 peptides per genome (Figure 2D). This variation indicates that raw peptide match counts are not directly comparable across genomes with different reference sizes. Together, these properties of the theoretical reference space explain why MetaUmbra retains shared peptides, adjusts them by degeneracy, and evaluates genome presence with formal statistical testing rather than raw peptide counts alone. A complementary observed-peptide diagnostic is shown in Supplementary Figure S2. When the SIHUMIx peptide set was matched against the full 4,744 genome UHGG representative background, matched peptides were broadly distributed across genomes and all background genomes retained at least one match, further illustrating why raw peptide matching alone is insufficient for genome presence inference.

### 3. Complete recovery of SIHUMIx with no additional significant genomes

The test panel contained 4,746 genomes, comprising the eight SIHUMIx genomes and 4,738 background genomes from the UHGG representative set after removal of six species-overlapping background genomes. Because the expected community contains only eight known members, this dataset tests whether MetaUmbra can recover the expected set while suppressing extra calls in a realistic gut microbiome reference space. MetaUmbra recovered all eight expected SIHUMIx members, with no additional calls among the 4,738 background genomes at both *q* ≤ 0.01 and *q* ≤0.05 (Figure 3A). Raw genome-level p-values for the expected members ranged from 2.0 χ 10^−300^ to 3.13 χ 10−8, and BH-adjusted q-values ranged from 3.164 χ 10^−297^ to 1.854 χ 10−5.

**Figure 3.**
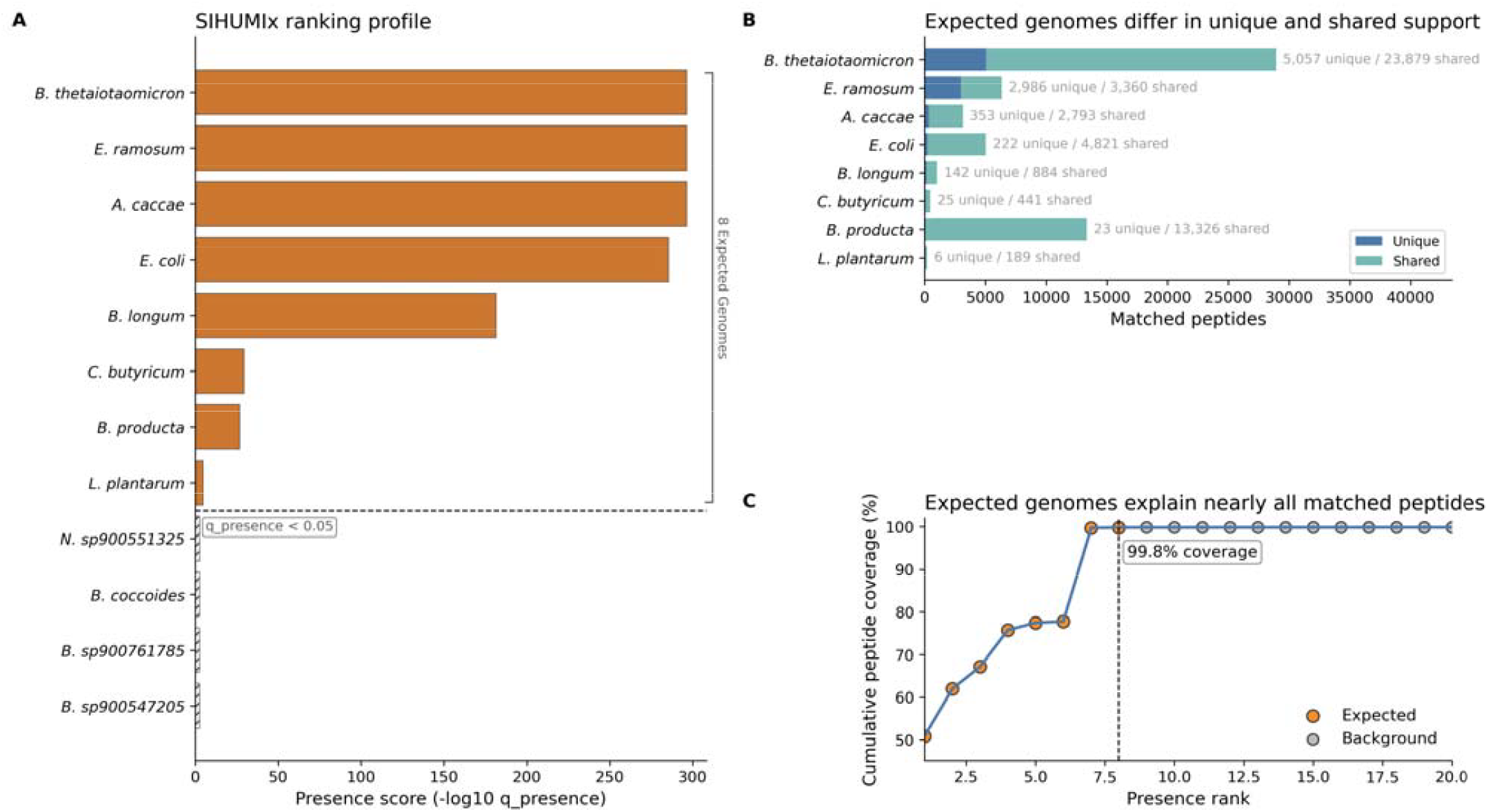
SIHUMIx benchmark results. (A) Presence scores, defined as, for the 12 highest-ranked candidates. The eight expected genomes occupy ranks 1–8; no additional genome is significant at or. The dashed line marks. (B) Unique and shared matched-peptide counts for the eight expected genomes, with both components annotated. (C) Cumulative matched-peptide coverage across the ranking, reaching 99.8% by rank 8.

The SIHUMIx ranking was also highly compact (Figure 3A, C). The top eight genomes accounted for 99.8% of cumulative matched-peptide coverage, separating the expected set from the UHGG background. The expected genomes occupied distinct unique and shared evidence regimes (Figure 3B): *Lactobacillus plantarum* had 6 unique and 189 shared matched peptides, and *Clostridium butyricum* had 25 unique and 441 shared peptides. *Erysipelatoclostridium ramosum* and *Bacteroides thetaiotaomicron* had substantially larger unique/shared counts of 2,986/3,360 and 5,057/23,879, respectively. Every expected genome retained unique support, while shared evidence described complementary aspects of its peptide profile. Full candidate-level results are provided in Supplementary Table S2.

All 4,746 SIHUMIx candidates had at least one matched peptide, yet only the eight expected members were significant. The 56,852 distinct matched peptide sequences spanned a broad degeneracy range (Figure 4A). The evidence profiles distinguished weak expected members from shared-rich background competitors (Figure 4B): *Lactobacillus plantarum* and *Clostridium butyricum* remained significant with only 6 and 25 unique peptides, respectively, whereas candidates supported exclusively by shared peptides received under the zero-unique rule. Stratum-level profiles further showed that expected genomes differed in the degree of ambiguity associated with their shared evidence (Figure 4C). These results illustrate why raw peptide matches alone are insufficient to establish genome-level support.

**Figure 4.**
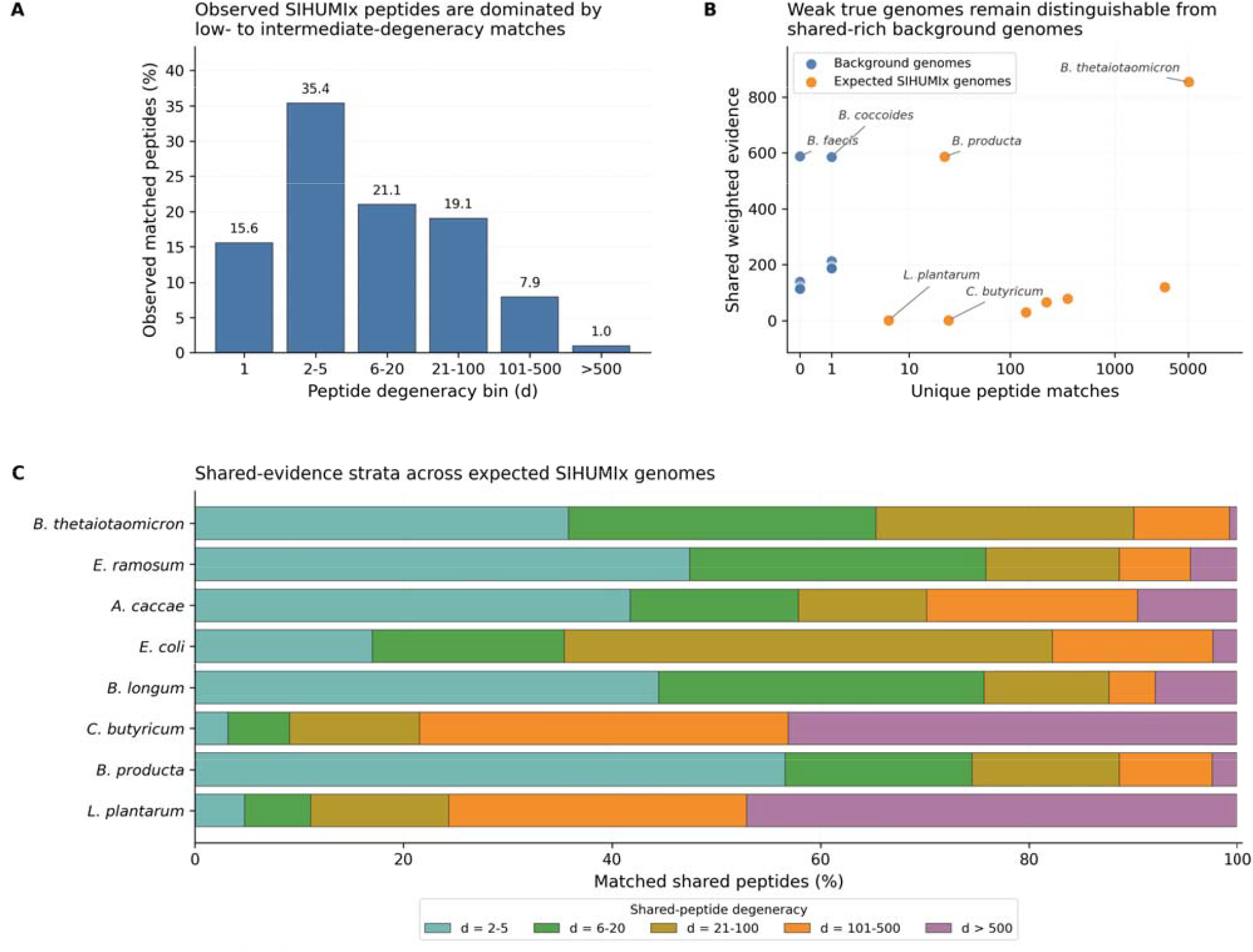
Evidence structure of the SIHUMIx benchmark. (A) Degeneracy distribution of 56,852 distinct matched peptide sequences across six candidate-genome multiplicity bins. (B) Unique matched-peptide counts and degeneracy-weighted shared evidence for the eight expected genomes and eight selected high-evidence background candidates. Unique counts use a coordinate so that zero-valued candidates remain visible. (C) Shared-assignment composition across five degeneracy strata for each expected genome; percentages use the shared assignments of that genome as the denominator.

## 4. Genome discrimination in Mix24X under expanded backgrounds

Mix24X served as the discrimination-focused benchmark because it contains distantly related species and difficult near-neighbor cases. Using the 24 individual-strain peptide sets, each scored against the exact 24 Mix24X genomes, MetaUmbra recovered the expected genome at in all 24 analyses. Twenty analyses had no additional call, whereas four had one additional call involving the *Bacillus cereus*/*Bacillus thuringiensis* and *Salmonella bongori*/*Shigella flexneri* pairs (Figure 5A). Thus, most individual-strain analyses recovered only their expected member, while closely related genomes remained a source of ambiguity.

**Figure 5.**
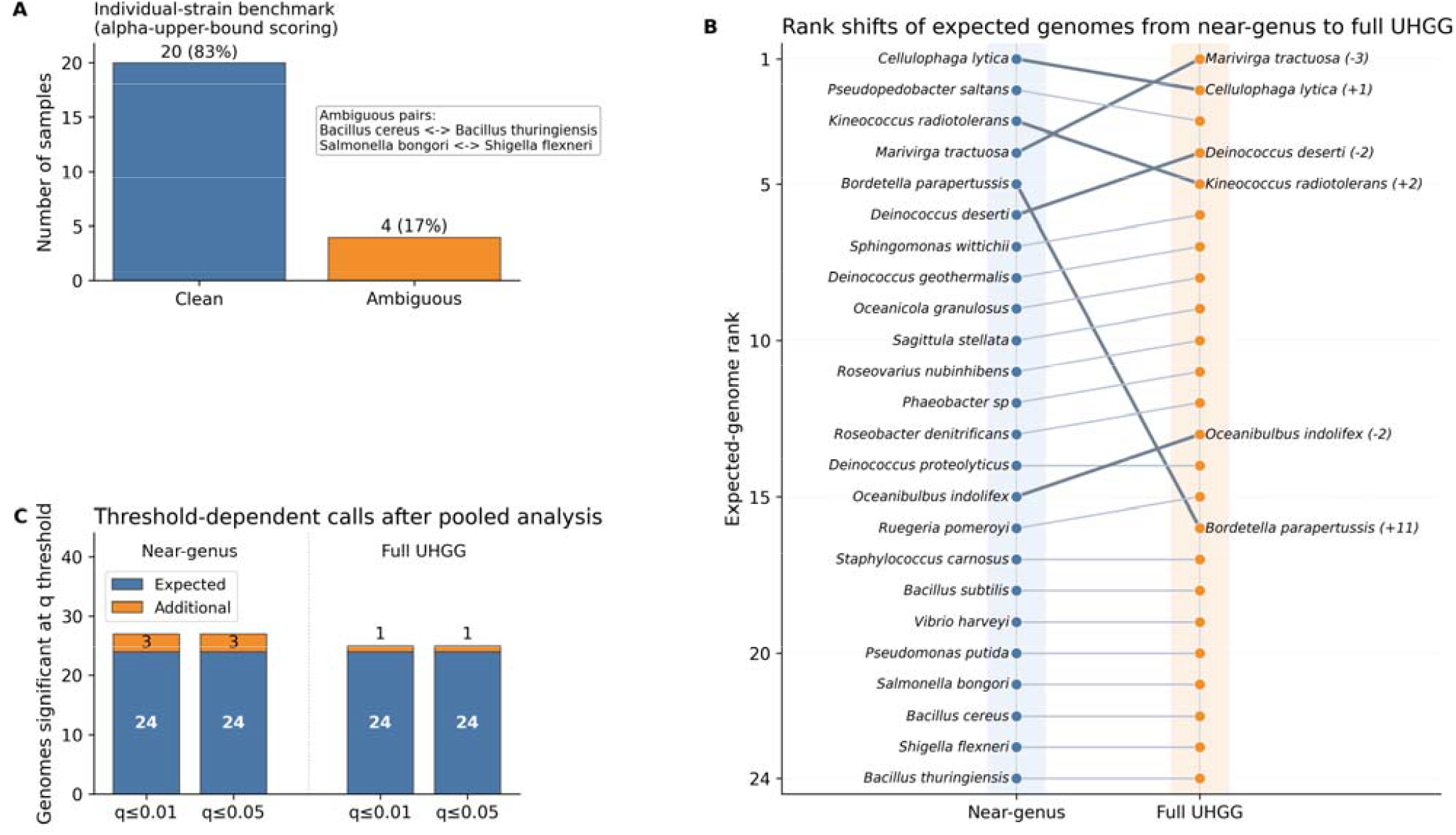
Mix24X benchmark results under background expansion. (A) Summary of 24 individual-strain analyses against the exact 24-genome panel at : every expected genome was recovered, 20 analyses had no additional call, and four had one closely related additional call. (B)Presence-rank shifts of the 24 expected genomes from the near-genus to the full-UHGG panel; all remain within ranks 1–24. (C) Numbers of expected and additional significant genomes at and for both pooled analyses.

We next analyzed a synthetic pooled peptide set generated by merging all 24 individual-strain peptide sets. The near-genus panel contained 73 genomes; the species-deduplicated full-UHGG panel contained 4,764 genomes after removal of four species-overlapping UHGG representatives. All 24 expected genomes remained within ranks 1–24 in both panels (Figure 5B). The largest full-minus-near rank shift was *Bordetella parapertussis*, from rank 5 to 16 (+11). *Marivirga tractuosa* shifted −3; *Kineococcus radiotolerans* shifted +2; *Deinococcus deserti* and *Oceanibulbus indolifex* each shifted −2. All other absolute shifts were at most one rank. These changes represent local reordering within the expected set rather than loss of its members.

Both pooled panels recovered all 24 expected genomes at and. The near-genus panel produced 3 additional calls among 49 non-expected candidates, whereas the full-UHGG panel produced 1 additional call among 4,740 non-expected candidates (Figure 5C). The former used alpha-upper-bound unique evidence, while the latter had adequate comparable background for the empirical-background model. Reference expansion therefore changed both the specificity of peptide support and the applicable unique-evidence model; it did not increase additional calls in these analyses. Full candidate-level results are provided in Supplementary Table S2.

To examine why expected-genome recovery was preserved despite reference expansion, we compared the significance-threshold response and peptide evidence structure. Across q-value thresholds from 0.001 to 0.2, the near-genus panel retained 3 additional calls, increasing to 9 at. The full-UHGG panel retained 1 additional call throughout (Figure 6A).

**Figure 6.**
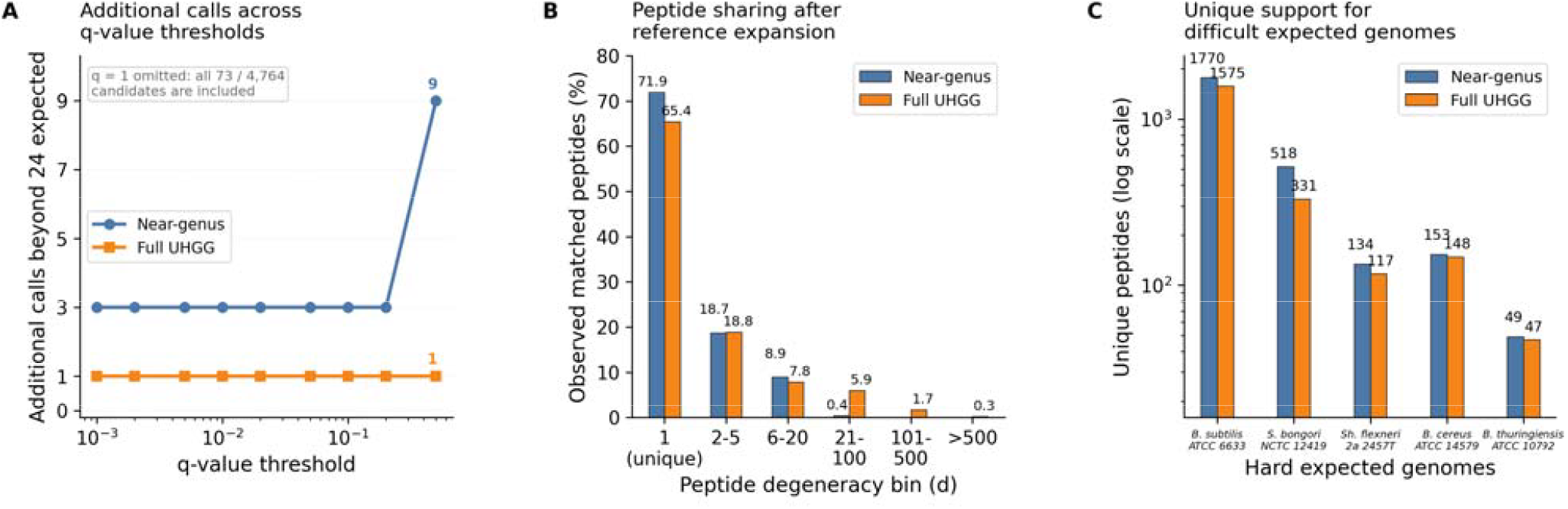
Evidence shifts in Mix24X under background expansion. (A) Additional calls beyond the 24 expected genomes across BH q-value thresholds from 0.001 to 0.5 in the near-genus and full-UHGG panels. The endpoint is omitted because it applies no significance filter. (B) Distribution of distinct matched peptide sequences across six candidate-genome multiplicity bins. (C) Unique-peptide support for five difficult expected genomes before and after expansion, shown on a logarithmic axi**Table 1. Acquisition, peptide-input fields, and analysis units for the primary datasets**.

Reference expansion redistributed peptides among unique and shared categories (Figure 6B). Across the 24 expected genomes, the median unique-peptide fraction decreased from 61.41% to 56.43%, and the median change was −83.5 unique assignments per genome. Unique support for difficult expected genomes declined but remained substantial in several cases: *Bacillus cereus* retained 148 of its 153 near-genus unique peptides, *Bacillus thuringiensis* retained 47 of 49, and *Shigella flexneri* retained 117 of 134 (Figure 6C). All 24 expected genomes remained significant. The effect of expanding the reference was therefore panel-dependent, with altered peptide specificity and local reordering but preserved expected-member recovery.

Additional external evaluations using two independent published mixture datasets are presented in Supplementary Figures S3 and S4.

## 5. Comparison with peptide based baseline tools

To place the revised MetaUmbra results in context, we compared them with retained Unipept 6.4.3 (6) and Peptonizer2000 (11) outputs for the same SIHUMIx and Mix24X peptide datasets (Figure 7). These tools address related taxonomic interpretation problems but operate at the taxon level rather than as explicit genome-level hypothesis tests within user-defined candidate panels. Because the external analyses were not rerun after revision of the MetaUmbra scoring procedure, the comparison was used descriptively to assess expected-member recovery and output compactness. Detailed Unipept and Peptonizer2000 outputs are provided in Supplementary Table S3.

**Figure 7.**
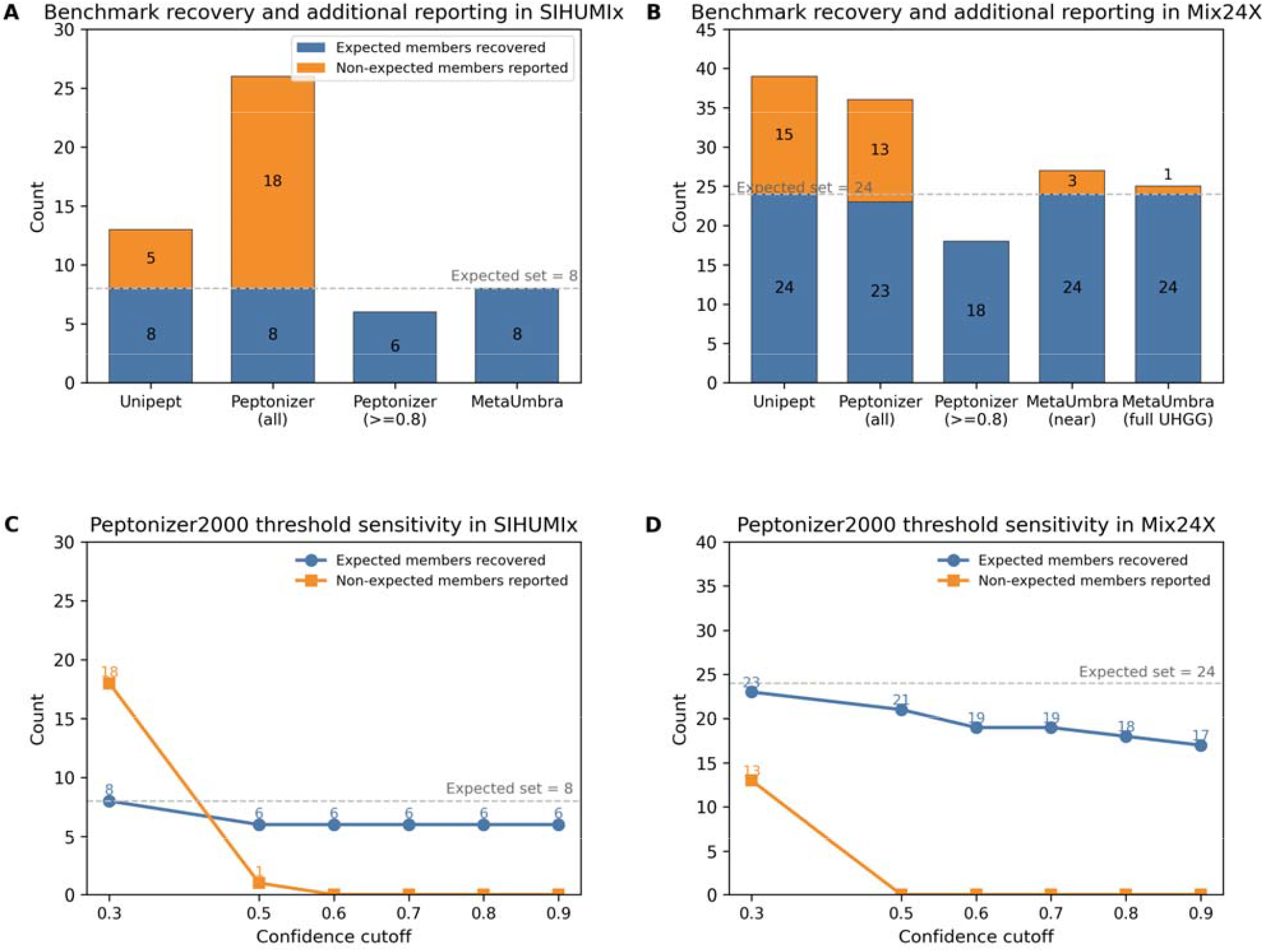
Comparison with peptide-based baseline tools. (A, B) Harmonized counts of expected and additional reported taxa or genomes for SIHUMIx and Mix24X. Updated synonyms and redundant aliases were collapsed before comparison. MetaUmbra uses the revised q ≤ 0.01 results, whereas Unipept and Peptonizer2000 values were retained from the previous external-tool analysis. (C, D) Peptonizer2000 results across confidence cutoffs from 0.3 to 0.9. Additional reports are defined relative to benchmark membership and should not be interpreted as estimates of BH false-discovery rate; taxon-level and genome-level outputs have different inferential targets.

In SIHUMIx, Unipept recovered all eight expected members and reported five additional taxa. Peptonizer2000 retained all eight expected members when all reported taxa were considered, but the harmonized output contained 26 taxa overall. At the recommended 0.8 confidence cutoff (11), Peptonizer2000 no longer retained *Clostridium butyricum* or *Lactiplantibacillus plantarum*. MetaUmbra retained all eight expected genomes with no additional significant calls at. Because MetaUmbra evaluates only genomes within the declared candidate panel, these outputs should be interpreted relative to the different reference scopes and inferential targets of the three methods.

In Mix24X, Unipept recovered all 24 expected species, with the harmonized output containing 39 taxa. Peptonizer2000 reported 36 taxa overall and recovered 23 of the 24 expected species. Increasing the confidence cutoff reduced the number of reported taxa but also removed several difficult expected members (Figure 7C, D). At the recommended 0.8 cutoff, *Bordetella parapertussis, Bacillus cereus, Bacillus thuringiensis, Pseudomonas putida, Shigella flexneri*, and *Cellulophaga lytica* were not retained. In comparison, MetaUmbra retained all 24 expected genomes in both expanded-panel analyses at, reporting 27 significant genomes in the near-genus panel and 25 in the full-UHGG panel. All expected genomes remained within ranks 1– 24.

### 6. Application of MetaUmbra to a hamster gut reconstructed genome panel

MetaUmbra was further applied to a hamster gut reconstructed genome panel comprising 706 annotated bacterial MAGs. Unlike SIHUMIx and Mix24X, this dataset does not define a closed expected set; therefore, the analysis was used to assess evidence structure in a dense real-data reference space rather than recovery of a predefined community. Using I/L-equivalent matching and the alpha-upper-bound unique-evidence model, 587 of 706 MAGs were significant at both q ≤0.01and q ≤0.05. Exact-sequence matching yielded 588 significant MAGs, with 578 shared between the two analyses (Jaccard index = 0.9682), indicating that the overall result was largely preserved while a small number of genome calls depended on I/L normalization.

The strongly supported MAGs displayed in Figure 8 included *Escherichia coli, Desulfovibrio* sp., and MAGs related to Bacteroidaceae, Muribaculaceae, Lachnospiraceae, and Bifidobacteriaceae. These lineages provide biological context for the peptide evidence observed in the reconstructed hamster panel, although their taxonomic plausibility does not independently validate genome presence.

**Figure 8.**
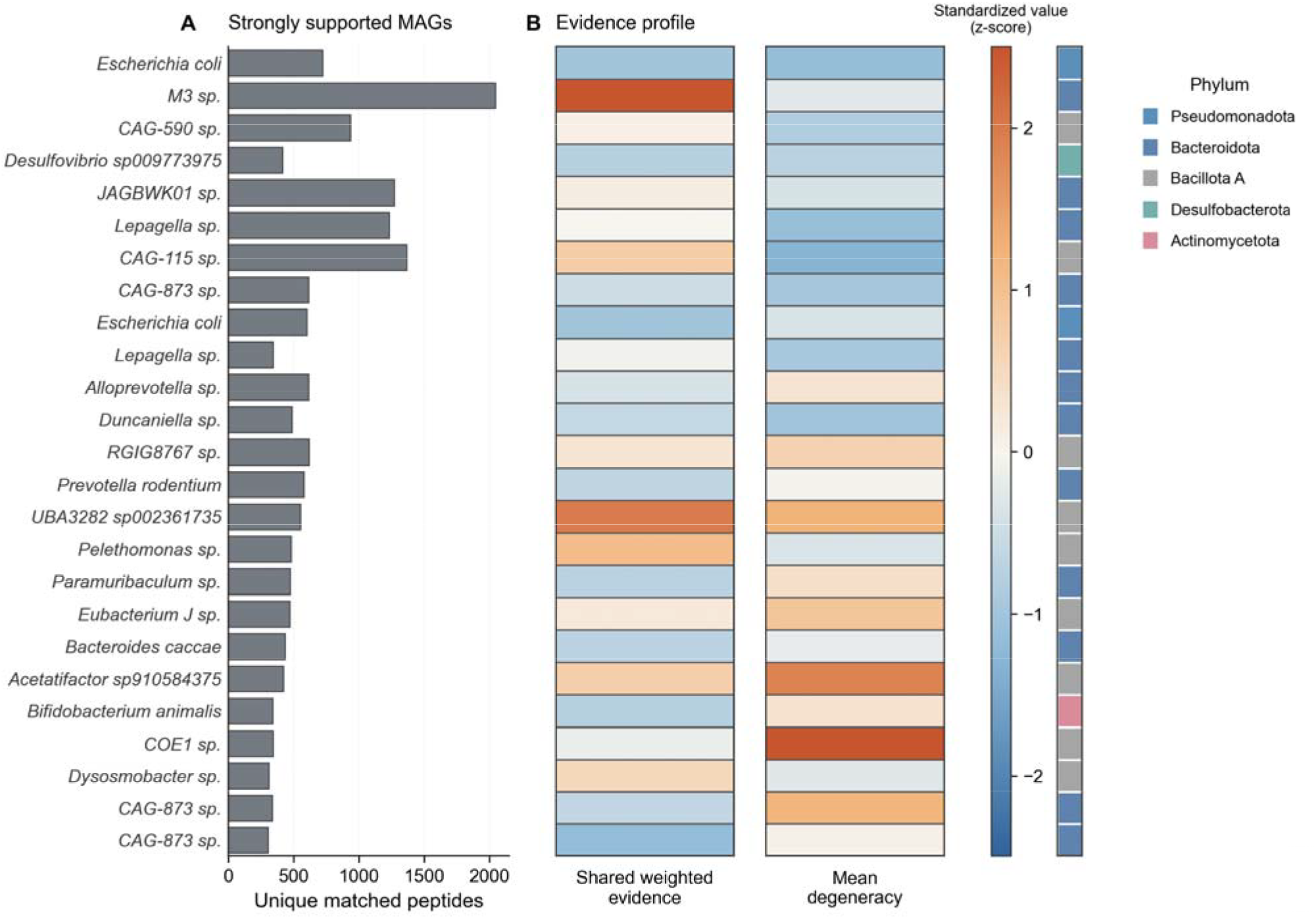
Evidence structure of strongly supported MAGs in the hamster reconstructed genome panel. (A) Unique matched-peptide counts for the displayed MAGs, ordered by the descriptive display score defined in the text. (B) Weighted shared evidence and mean peptide degeneracy for the same MAGs. Heatmap values are column-wise z-scores, and the annotation strip indicates phylum.

Figure 8 summarizes complementary evidence profiles among strongly supported MAGs. Because several candidates reached the same minimum formal q-value, formal significance alone could not provide a meaningful ordering within this tied group. We therefore defined a descriptive display score for visualization, calculated as the sum of the standardized log-transformed unique-peptide count and the standardized shared-evidence significance. The resulting profiles showed substantial unique-peptide support together with varying levels of weighted shared evidence and peptide degeneracy.

### 7. Robustness of genome calls and computational performance

Following the benchmark and hamster analyses, we evaluated the robustness of MetaUmbra ca ls by examining unique-peptide support, sensitivity to analysis settings and reference composition, and computational performance.

We first asked whether significant genome calls were being supported by only a very small number of unique peptides. No significant candidate was supported by zero or one observed unique peptide at either q-value threshold. Using 1-3 unique peptides as a descriptive low-support category, such records accounted for 0/8 significant SIHUMIx candidates, 2/27 pooled Mix24X near-genus candidates, 0/25 pooled full-UHGG candidates, and 1/28 candidates across the 24 individual Mix24X analyses. All three low-support benchmark records were additional calls and each had three unique peptides. In the hamster panel, 71/587 significant MAGs in the I/L-equivalent analysis and 71/588 in the exact-sequence analysis fell within this low-support category. Full support-count distributions are provided in Supplementary Table S7.

We next tested whether the formal call sets were sensitive to stochastic variation or the choice of peptide-degeneracy bins. Across all 27 combinations of three benchmark panels, three binning schemes, and three random seeds, the significant call sets were identical to the corresponding default analyses at both reporting thresholds (Jaccard index = 1). Thus, the benchmark-level conclusions were stable to these implementation choices.

We then examined a different source of uncertainty: incompleteness of the candidate reference panel. In leave-one-reference-out analyses, all remaining expected genomes were retained, including 7/7 SIHUMIx members and 23/23 Mix24X members in each corresponding omission scenario. However, removing an expected reference also produced additional genome calls, indicating that reference composition can alter the specificity of existing peptide evidence. At q ≤ 0.01, 73 SIHUMIx and 59 Mix24X candidate-omission events became newly significant.

Examination of these newly significant calls showed a consistent mechanism. Their total matched-peptide counts remained unchanged, while peptides previously shared with the omitted reference became panel-unique to another candidate. Across the 132 newly significant events, panel-unique counts increased and shared counts decreased correspondingly, and 117 candidates had no panel-unique peptide in the baseline analysis. Sequence-level mapping confirmed this interpretation: 9,512 SIHUMIx and 1,334 Mix24X peptide-event records gained panel uniqueness after omission, and each sequence had originally matched exactly the newly called candidate and the omitted reference. Supplementary Figure S5 summarizes these changes, and Supplementary Tables S5 and S6 provide the paired results and sequence-level mappings.

Finally, we evaluated the computational requirements of reference scoring and preparation. Component scoring required 43.6 s and 2.35 GiB peak process-tree memory for the 73-genome near-genus panel, compared with approximately 18 min and 47 GiB for the 4,764-genome full-UHGG and 4,746-genome SIHUMIx panels. Reference digestion was benchmarked separately on a laptop using 16 worker processes: digestion of 100, 200, and 500 locally stored UHGP genome FASTA files required 69.7, 139.5, and 345.9 s, respectively, with peak process-tree memory remaining below 0.44 GiB. Detailed hardware specifications, input and output sizes, and conditional extrapolations are provided in Supplementary Table S8.

## Discussion

A central challenge in metaproteomics is that peptide evidence must be interpreted in the presence of extensive sharing across related organisms. This makes it difficult to determine whether a specific candidate genome is truly supported by the observed peptide set. MetaUmbra was developed to address this problem by assigning each candidate genome a formal genome-level significance value for presence. Across the SIHUMIx and Mix24X benchmarks, MetaUmbra recovered all expected genomes, while additional calls remained dependent on reference composition. In the hamster MAG application, it further provided interpretable complementary evidence profiles in a dense real-data reference space.

A practical but limited strategy in peptide-based taxonomic analysis is to prioritize candidate taxa on the basis of fixed unique-peptide counts, for example by retaining only taxa supported by more than three unique peptides. However, such rules are inherently sensitive to the size and composition of the reference background. A peptide that appears unique in a smaller or sparser database may no longer remain unique after background expansion or after inclusion of more closely related genomes. Fixed unique-peptide thresholds therefore do not provide a stable basis for presence inference across different reference spaces. MetaUmbra addresses this limitation by jointly evaluating unique and shared peptide evidence within a formal statistical framework. Shared peptides are retained and weighted according to their degeneracy across the reference panel, allowing ambiguous peptide evidence to contribute without being treated as equivalent to genome-unique support. The testing procedure further constrains genome-level inference so that shared evidence alone is insufficient to establish a formal presence call. This design avoids reliance on arbitrary unique-peptide count thresholds while explicitly incorporating the information carried by shared peptides.

The comparison with Unipept and Peptonizer2000 further defines the methodological position of MetaUmbra. Unipept uses lowest-common-ancestor assignment for peptide-based taxonomic annotation, whereas Peptonizer2000 assigns taxon-level confidence scores that can be used for candidate prioritization. MetaUmbra instead provides formal genome-level significance values, including raw p-values and BH-adjusted q-values, within a user-defined candidate panel. These outputs address related but distinct inference tasks and should not be interpreted as directly equivalent measures. In the benchmark datasets, MetaUmbra retained all expected genomes while maintaining compact significant sets, whereas the retained taxa from confidence-based approaches depended more strongly on the reporting threshold. For MetaUmbra, the choice of q-value threshold is distinct from the handling of reference complexity, which is incorporated during evidence evaluation. The final threshold instead reflects reporting stringency; here, *q* ≤ 0.01 was used as the primary threshold and *q* ≤ 0.05 for secondary review.

The hamster application extends this interpretation to a complex real-data setting. Unlike SIHUMIx and Mix24X, it did not involve a closed expected set and therefore should not be interpreted as a recovery benchmark. The displayed MAGs combined substantial unique-peptide support with varied shared evidence, whereas the large number of significant MAGs should be interpreted in the context of the complex and related candidate panel. Thus, this analysis illustrates how MetaUmbra organizes genome-level peptide evidence in a dense reference space, while individual calls remain conditional on panel composition.

MetaUmbra inference is conditional on the composition and completeness of the reference panel. Leave-one-reference-out analyses showed that removing an expected reference can convert previously shared peptides into panel-unique evidence for another candidate and generate additional calls. Panel-unique support should therefore not be interpreted as universally species-specific, and significant genomes should be considered together with plausible close competitors and reference completeness. Discrimination among very closely related genomes remains limited by extensive peptide sharing, although iterative refinement from broad panels to more densely sampled species-or strain-level references may improve resolution. Additional limitations include dependence on upstream peptide-identification quality, score calibration, and digestion assumptions; fragment-level dependence in DIA data is not explicitly modeled. MetaUmbra is intended as a genome-presence module within broader metaproteomics workflows and does not itself estimate abundance or perform differential or functional analysis.

## Conclusions

MetaUmbra converts metaproteomic peptide evidence into genome-level significance values for candidate genome presence. By integrating unique-peptide support with ambiguity-aware statistical evaluation of shared peptides, it provides a formal framework for genome-level presence inference in peptide-centric metaproteomics. In SIHUMIx and Mix24X, MetaUmbra recovered all expected genomes, while additional significant calls remained dependent on reference composition. In the hamster reconstructed genome panel, it also provided interpretable complementary evidence profiles in a dense real-data setting. Although UHGG was used here as a representative human gut background, the same framework is applicable to other environments, including marine, soil, and other host-associated microbiomes. MetaUmbra can also be applied iteratively to refine genome-level inference within more closely related reference groups.

## Supporting information

MetaUmbra_supplementary_materials

## Abbreviations

ATCC: American Type Culture Collection
BH: Benjamini–Hochberg
DDA: data-dependent acquisition
DIA: data-independent acquisition
FDR: false discovery rate
MAG: metagenome-assembled genome
Mix24X: 24-species single-strain benchmark dataset
PEP: posterior error probability
PRIDE: Proteomics Identifications Database
SIHUMIx: simplified human intestinal microbiota
SPEED: sample preparation by easy extraction and digestion
TPM: transcripts per million
UHGG: Unified Human Gastrointestinal Genome

## Acknowledgment

This work was supported by the Natural Sciences and Engineering Research Council of Canada (NSERC) through grant RGPIN-03905-2018 to D.F.

## Data Availability

Primary benchmark data analyzed in this study are available from PRIDE under PXD023217 (SIHUMIx) and PXD005728 (Mix24X). The hamster metagenomic reference resource is available at Zenodo record 13909289, and the matched metaproteomics dataset is available under PXD057158. Supplementary external evaluations used PXD057701 and PXD011189. Processed peptide inputs are provided in Supplementary Table S4, and revised candidate-level statistical results and evidence counts are provided in Supplementary Table S2. Reference-omission analyses and sequence-level mappings are provided in Supplementary Tables S5 and S6, respectively. Unique-support summaries and computational benchmarks are provided in Supplementary Tables S7 and S8.

MetaUmbra can be used as a standalone tool through a graphical user interface, command-line interface, or function-based workflow. It has also been integrated into the peptide annotation workflow of MetaX v2.0 (33). Source code is available from the MetaUmbra GitHub repository (https://github.com/byemaxx/MetaUmbra).

## Author Contributions

Q.W.: Conceptualization, methodology, software, validation, formal analysis, investigation, data curation, visualization, writing – original draft, and writing – review and editing. Z.N.: Conceptualization, methodology, supervision, and writing – review and editing. A.Z.: Resources, data curation, investigation, and writing – review and editing. K.C.: Resources and writing – review and editing. D.F.: Conceptualization, supervision, project administration, funding acquisition, resources, and writing – review and editing.

## Declaration of Generative AI and AI-assisted Technologies in the Writing Process

During the preparation of this work, the authors used ChatGPT (OpenAI) to assist with English language editing and to improve clarity and readability. The tool was not used to generate scientific hypotheses, perform data analysis, interpret results, or draw conclusions. After using this tool, the authors reviewed and edited the manuscript as needed and take full responsibility for the content of the published article.

