## Supplementary material for "MetaUmbra: Statistically Controlled Genome-Level Presence Inference from Metaproteomic Peptides": MetaUmbra_supplementary_materials

Supplementary Results
Graphical user interface
To support routine use, MetaUmbra was implemented with a graphical user interface in addition to command-line and function-based workflows. The interface provides entry points for genome presence scoring and FASTA digestion. In the genome presence scoring tab, users can select an observed peptide table, an optional genome-lineage annotation table, an output result file, and one or more genome digest directories; users can also define peptide sequence and score columns and access additional runtime options for Monte Carlo scoring and matched-peptide caching (Supplementary Fig. S1). This interface is intended to make the workflow accessible for users who need genome-level presence inference from peptide lists without directly editing scripts.
SIHUMIx matched-peptide distribution across the UHGG background
To visualize nonspecific peptide matching before statistical scoring, Supplementary Figure S2 summarizes all 4,746 candidates in the retained SIHUMIx result table, including the UHGG representative background and expected-community references. Matched peptide counts were broadly distributed, with a first quartile of 111, a median of 164, and a third quartile of 367 matched peptides per candidate. All 4,746 candidates had at least one match. Thus, a simple one-match threshold would nominate the entire candidate panel. This diagnostic motivates evaluating unique and degeneracy-weighted shared evidence against genome-specific null expectations rather than using raw peptide occurrence alone.
External mixture datasets and reference-background design. Two independent published mixture datasets were included as supplementary evaluations. For both datasets, the source-study reference taxa were evaluated together with the UHGG representative genome background using the same background-expansion strategy applied to the primary benchmarks.
Six-strain culture-mixture dataset
The first supplementary dataset was derived from the metagenome-informed metaproteomics study by Valdés-Mas et al. (1). We used the source study's “taxonomic profiling by proteomic signature” experiment. In that experiment, cultures of Staphylococcus aureus ATCC 12600, Klebsiella oxytoca ATCC 13182, Micrococcus luteus DSM 20030, Escherichia coli K-12 ATCC 700926, Enterococcus faecalis ATCC 29212, and Bacillus subtilis strain 168 ATCC 23857 were grown aerobically at 37 °C in Mueller Hinton broth. Optical density was equalized to 1.0, and isovolumic culture mixtures were generated. The source study also included 14 absent gut bacterial species in its protein reference database to simulate real-world conditions and allow erroneous taxonomic identification to occur (1).
In the MetaUmbra evaluation, the six expected reference taxa were embedded in the UHGG representative genome background. MetaUmbra ranked the six expected taxa in the top six positions, and all six were significant at q ≤ 0.01 (Supplementary Fig. S3). No non-reference genome was significant at q ≤ 0.05. The first non-reference entry was MGYG000001694, a UHGG Enterococcus faecalis representative, at rank 7. This background entry had extensive peptide matching but only one genome-unique peptide and was not significant after multiple-testing correction (p = 1.02 × 10−3; q = 0.378). This pattern indicates that MetaUmbra retained the expected source-study taxa while suppressing a shared-rich closely related background entry.
The six-strain result illustrates the contrast between shared-rich background matching and statistically supported reference taxa, providing an additional evaluation of MetaUmbra beyond the primary SIHUMIx and Mix24X benchmarks.
ATCC 20-strain microbiome standard
The second supplementary dataset was derived from the SPEED sample preparation study by Doellinger et al. (2). The original study used the commercial 20 Strain Even Mix Whole Cell Material microbiome standard (ATCC MSA-2002). The lyophilized standard was resuspended in PBS, divided into aliquots, pelleted, and then processed for proteomic analysis. In the source study, the microbiome standard was prepared in triplicate using SPEED, STrap, and iST and analyzed by 4 h single-shot LC-MS/MS (2). Here, the SPEED-derived ATCC 20-strain peptide evidence was used for the supplementary MetaUmbra evaluation.
In the MetaUmbra evaluation, the ATCC reference taxa were evaluated together with the UHGG representative genome background. MetaUmbra placed all 20 ATCC reference taxa within ranks 1–20 (Supplementary Fig. S4). Nineteen of the 20 reference taxa were significant at q ≤ 0.05, and no non-reference genome was significant at the same threshold. The remaining reference taxon, Propionibacterium acnes (source-study name retained), was ranked at position 20 but did not pass the q ≤ 0.05 threshold (p = 5.19 × 10−4; q = 0.123). This taxon had limited genome-specific support in the processed peptide set, with 123 matched peptides and three unique peptides. The first non-reference entry, MGYG000002548, was ranked at position 21 and was far below the significance threshold (q = 0.928).
The ATCC standard therefore represents a more complex mixture than the six-strain culture-mixture dataset and includes a low-support boundary case. The failure of Propionibacterium acnes to pass the adjusted threshold should be interpreted as a sensitivity limitation for a reference taxon with limited unique-peptide evidence rather than as nonspecific expansion of the call set, because no UHGG background genome was significant. This behavior is consistent with the intended use of MetaUmbra, in which candidate genomes are evaluated within the supplied reference collection and retained as significant only when their evidence remains sufficient after multiple-testing adjustment.
Supplementary Methods: adaptive background and peptide-input processing
For empirical-background scoring, let N be the number of genomes with at least one observed peptide match. At exclusion fraction f, the number removed from the background was min(ceil(Nf), N − 1), preserving at least one background genome when N > 0. Genomes were ordered by decreasing unique-peptide count, decreasing unique weighted evidence, decreasing total weighted evidence, and increasing genome identifier. After exclusion, quantile-bin boundaries were recalculated from theoretical panel-unique opportunity in the retained background. The requested bin count was limited by the structural partition (at most eight bins), floor(background size/50), and the number of distinct opportunity values, with at least one bin. Duplicate quantile boundaries were removed; a degenerate partition or binning failure produced a single bin. Each candidate was compared with retained genomes in its opportunity bin, excluding itself. An empty comparison bin fell back to the whole retained background excluding the candidate. The 95th percentile used linear interpolation; an empty background used a threshold of zero. The nonnegative excess of the observed unique count over this threshold entered the α-excess statistic defined in Experimental Procedures section 3.
All reported primary analysis families used all-samples mode, including the individually selected Mix24X strain inputs. When empirical-background scoring was active, f began at 0.10. After each update of the unique statistics, the unique- and shared-evidence p-values were evaluated using a two-component Simes intersection test followed by closed testing of the unique-evidence component and BH adjustment across the N matched genomes. The fraction with intermediate q ≤ 0.20 was clipped to [0.05, 0.20] to obtain the next f. The update was accepted before checking convergence (absolute change <0.01); at most five updates were performed. A final recomputation used the last updated fraction, even if the iteration limit was reached. Shared Monte Carlo refinement and final BH adjustment followed, without another background update. The software also provides a grouped-unit profile (initial fraction 0.05, bounds [0.02, 0.20], three updates), but that profile was not used for these analyses. The alpha-upper-bound analyses did not use adaptive empirical background subtraction in their formal unique p-values. These implementation settings describe the retained analyses and are not independent validation of the component p-values.
The resolved input fields, rather than unfulfilled column requests in generic configurations, determine preprocessing. SIHUMIx used Sequence, Score, PEP (≤0.05), and Reverse (entries marked + removed); Intensity supplied the presence filter. The saved SIHUMIx run loaded 66,178 rows, retained 66,013 after decoy removal, and used 60,529 distinct sequences after the remaining preprocessing. Mix24X used Sample, Sequence and Score, with no resolved peptide-error or decoy column; no additional error/decoy filter was applied by MetaUmbra to these published filtered identifications. Hamster scoring used the reduced table containing Sequence, score and Q.Value, with Q.Value ≤0.05 and no decoy column. The score field contained the retained Evidence-derived relative support; PEP from the upstream report was not a column in this reduced scoring input. Mix24X and hamster inputs without an intensity column used synthetic presence values. For all three datasets, sequence-level maximum scores were min–max scaled within each scoring run as described in Experimental Procedures section 3. The fixed unique-evidence error bound α = 0.05 was a separate model parameter, not a transformation of Score, Evidence, PEP or Q.Value.
The primary analysis used a two-component Simes intersection test followed by closed testing of the unique-evidence component. For genomes without an observed unique peptide, the final genome-level p-value was set to 1. Candidate-level outputs include the unique- and shared-evidence p-values, the Simes intersection p-value, the final genome-level p-value, and the BH-adjusted q-value. Reference-omission and bin/seed sensitivity analyses were rerun using MetaUmbra 1.5.0 as described below.
Reference omission and additional genome calls
Reference omission and bin/seed sensitivity were rerun with MetaUmbra 1.5.0 using the retained manuscript settings. Three panels (SIHUMIx, pooled Mix24X near-genus and pooled Mix24X full-UHGG), three bin schemes and seeds 1, 7 and 42 yielded 27 runs. The bin edges were [1, 5, 20, 100, 500] (default), [1, 20, 100, 500] (coarse) and [1, 2, 5, 10, 20, 50, 100, 500] (fine). The 32 omission runs used default bins and seed 1. Each run used 500 initial Monte Carlo iterations and 20,000 refined iterations for the near-genus panel or 200,000 for the two full-background panels. The identified peptide inputs and other scoring settings were unchanged. One expected genome was removed before degeneracy, opportunity and scoring were recalculated. This baseline retained the original panel selection and exclusion rules and did not denote an exhaustive reference collection.
At q ≤ 0.01, the eight SIHUMIx scenarios retained 7/7 expected genomes, with 1–44 additional calls; the 24 Mix24X near-genus scenarios retained 23/23, with 3–16 additional calls. At q ≤ 0.05, recovery was unchanged and the ranges were 1–50 and 3–16. All 27 bin/seed configurations retained the corresponding baseline call sets at both thresholds. At q ≤ 0.01, the 73 SIHUMIx extra-call events were all newly significant; the 131 Mix24X events comprised 72 baseline and 59 new events. At q ≤ 0.05, there were 83 new events in each benchmark. All 132 and 166 new events at the respective thresholds had unchanged matched counts, increased unique counts and equal decreases in shared counts; 117 and 142 began with no unique sequences. The final combined p-value was twice the unique p-value in all these events. Shared p-values decreased in 21/132 and 27/166 events, respectively. These nested threshold summaries are not independent samples.
The three baseline Mix24X near-genus additional genomes were MGYG000001279 (Pseudomonas_E fulva; 3 unique sequences; q = 6.7593 × 10⁻⁴), MGYG000002331 (Vibrio parahaemolyticus; 13; q = 7.1289 × 10⁻¹⁷) and MGYG000002366 (Salmonella enterica; 3; q = 3.5096 × 10⁻⁴). Only MGYG000002331 was an additional call in the full-UHGG baseline at q ≤ 0.01 (q = 4.6523 × 10⁻¹⁵). Bacteroides thetaiotaomicron omission yielded 44 additional calls, including 25 Bacteroides, four Phocaeicola and two Parabacteroides labels; five references lacked a resolved species label. Staphylococcus carnosus omission yielded 16 additional calls: three baseline calls and 13 new calls, including 11 Staphylococcus labels, Staphylococcus_A lentus and Bacillus_J andreraoultii. These annotations describe reference labels, not measured phylogenetic distances; the near-genus panel was already selected using taxonomic relationships.
Supplementary Table S5. Paired additional reference-genome calls after omission. The tab-separated table contains 442 records indexed by omission scenario, q threshold and genome ID. It reports the omitted member, reference label and lineage, whether the call was newly significant, baseline and omission matched/unique/shared counts, theoretical opportunity, empirical excess, component and combined raw p-values, and BH-adjusted q-values. Counts refer to distinct I/L-equivalent sequences in the pooled analysis unit. The same_genus_label field compares the first word of each annotation exactly, without resolving synonyms or genus suffixes, and is not a phylogenetic measure. SIHUMIx lineages agreed with the saved scoring diagnostics; Mix24X near-genus names were assigned post hoc. Missing species labels remain UNRESOLVED. Equal unique gains and shared losses follow from the count partition when matched counts are unchanged; they are not independent sequence-tracing evidence. Sequence-level reference ownership was verified separately and is reported in Supplementary Table S6. Biological-source attribution, ANI, contamination and reference-annotation validation were not performed.
Sequence-level reference mapping. We constructed a peptide-to-reference ownership map from the retained matched-peptide caches, whose sequence and digest contents were unchanged across software versions. For each newly significant candidate at q ≤ 0.01 or q ≤ 0.05, we enumerated baseline and post-omission unique sequence sets and recorded the complete reference owner set of each post-omission unique sequence. At q ≤ 0.01, all 9,512 SIHUMIx and 1,334 Mix24X gained peptide-event records had exactly the candidate and omitted reference as baseline owners; omission left a single owner. At q ≤ 0.05, the corresponding counts were 9,542 and 1,373 across 83 newly significant events in each benchmark. These threshold sets overlap. Reconstructed matched, unique and shared counts matched all 44,507 fresh genome-result rows. This confirms the reference-mapping change and does not identify the biological sources of the measured peptides.
Supplementary Table S6. Sequence-level reference ownership for newly significant leave-out calls. The tab-separated table contains 10,939 peptide-event records for 166 newly significant events at q ≤ 0.05: 10,915 gained unique and 24 retained unique records. A separate field marks the 132 events newly significant at q ≤ 0.01. Each record includes the omission scenario, candidate and omitted reference annotations, normalized peptide string, baseline and post-omission reference owner sets and degeneracy, classification as gained or retained unique, and event-level raw p-values and BH-adjusted q-values. J represents I/L equivalence. Event-level p/q values are repeated for sequence lookup and are not peptide-level significance values. A record is a sequence within a genome/omission event, not an independent biological observation. The table verifies reference matches; ANI, biological-source attribution, contamination and reference-annotation validation were not performed.
Supplementary Table S7. Observed unique-peptide support among significant candidate records. The TSV provides dataset and family summaries separately at q ≤ 0.01 and q ≤ 0.05, with counts and denominators for 0, 1, 2, 3 and at least 4 unique peptides. The displayed counts below apply to both thresholds, whose call sets were identical. Counts refer to distinct I/L-equivalent sequences except for the exact-matching hamster sensitivity analysis. Expected/additional labels use the declared benchmark genome set; unknown denotes the absence of a closed truth set. Zero-denominator fractions are left blank. Dataset and family rows are alternative summaries and must not be added together. Candidate-level component p-values, genome-level p-values, BH-adjusted q-values, and unique-peptide counts are provided in Supplementary Table S2.
Analysis / truth status	Calls	U=0	U=1	U=2	U=3	U≥4
SIHUMIx; expected	8	0	0	0	0	8
SIHUMIx; additional	0	0	0	0	0	0
Mix24X near-genus; expected	24	0	0	0	0	24
Mix24X near-genus; additional	3	0	0	0	2	1
Mix24X full-UHGG; expected	24	0	0	0	0	24
Mix24X full-UHGG; additional	1	0	0	0	0	1
Mix24X individual (24 analyses); expected	24	0	0	0	0	24
Mix24X individual (24 analyses); additional	4	0	0	0	1	3
Hamster I/L; unknown	587	0	0	31	40	516
Hamster exact; unknown	588	0	0	37	34	517

Computational benchmarks were performed on a Windows 11 Pro version 25H2 laptop with a 13th Gen Intel(R) Core(TM) i9-13900H processor (2.60 GHz) and 32.0 GB installed RAM (31.7 GB usable). Component scoring required 43.6 s and 2.35 GiB peak process-tree resident memory for the 73-genome Mix24X near-genus panel, 1,085.7 s and 46.94 GiB for the 4,764-genome full-UHGG panel, and 1,107.7 s and 46.69 GiB for the 4,746-genome SIHUMIx panel. These measurements describe genome-level evidence computation and exclude upstream peptide identification and one-time reference digestion. No GPU was used.
Supplementary Table S8. Local first-time digestion resource measurements and conditional extrapolations. The TSV separates measured records from estimated records. Peptide rows are protein–peptide associations before panel-wide I/L normalization or deduplication, not counts of globally distinct peptides. Resident memory is the sampled sum across the process tree and can count shared pages more than once. Timings include process startup and TSV writing but exclude network transfer, staging, validation and scoring. The three sizes are nested subsets, each measured once with 16 workers; no replicate-based error bars are available. Estimated time and disk-size ranges use the smallest and largest observed per-genome rates for the 16-worker configuration; they are planning ranges under comparable protein content, not statistical confidence intervals. Memory is reported only for measured runs.
Genomes	Workers	Proteins	Residues (million)	Wall time (s)	Peak RSS (GiB)	TSV (GiB)
100	16	218,455	70.73	69.7	0.429	0.448
200	16	427,318	137.40	139.5	0.429	0.873
500	16	1,087,574	349.62	345.9	0.436	2.234

To measure one-time reference preparation separately from scoring, we randomly sampled 500 protein FASTA files from an available 4,516-genome UHGP collection (seed 42), copied them from network storage to local disk with SHA-256 verification, and selected nested 100-, 200- and 500-genome subsets using seed 42. Each subset was digested once with 16 worker processes on a laptop running Windows 11 Pro version 25H2 with a 13th Gen Intel(R) Core(TM) i9-13900H processor (2.60 GHz) and 32.0 GB installed RAM (31.7 GB usable). Every run used a fresh output directory, MetaUmbra 1.5.0, RPG 2.0.5, RPG Trypsin enzyme ID 42, up to two missed cleavages and peptide lengths of 7–30 residues. These runs form a separate implementation-cost benchmark, not a reconstruction of the historical reference digestion. Wall time included process startup, digestion and TSV writing; peak summed process-tree resident memory was sampled every 0.1 s. Network transfer, input staging, output validation, peptide identification and genome scoring were excluded. No GPU was requested and the operating-system file cache was not flushed.
For planning only, multiplying the minimum and maximum observed seconds per genome by 4,516 gives approximately 52.1–52.5 min with 16 worker processes on this laptop. This range is a conditional extrapolation, not a measured full-library runtime or a confidence interval; it assumes similar protein content per genome and sustained local throughput. The available 4,516-genome source collection is distinct from the historical 4,744-genome manuscript background. Peak memory was not extrapolated linearly with genome count.
MetaUmbra does not require a specific precomputed reference library. Users can generate reference libraries from genome-specific protein FASTA files appropriate to their study. For gut microbiome applications, publicly available MGnify resources such as UHGG provide a practical starting point for reference construction and testing. Once generated, genome-specific peptide reference tables can be reused when the genome versions and digestion settings remain unchanged; panel-dependent peptide normalization, matching, and degeneracy are recalculated during scoring.
Supplementary Figures


Supplementary Figure S1. MetaUmbra graphical user interface. The screenshot shows the genome presence scoring tab, including input selection for observed peptide tables, optional genome lineage annotation, output files, genome digest directories, column mapping, and access to optional runtime settings. The interface also provides access to FASTA digestion and peptide-table import workflows.


Supplementary Figure S2. Distribution of SIHUMIx observed peptide matches across the expanded candidate panel. (A) Histogram of matched peptides per candidate on a log scale. The median was 164, with an interquartile range of 111 to 367. (B) Ranked matched peptide counts showing that all 4,746 candidates had at least one matched peptide. The one-match threshold is shown as a dashed line.


Supplementary Figure S3. MetaUmbra evaluation of the six-strain culture-mixture dataset. (A) Presence-score ranking of the top candidate entries after evaluating the six source-study reference taxa together with the UHGG representative genome background. The six expected taxa occupied ranks 1–6 and were significant at q ≤ 0.05. (B) Screening context showing the number of candidate entries, reference taxa, reference taxa called at q ≤ 0.05, and non-reference calls at q ≤ 0.05.


Supplementary Figure S4. MetaUmbra evaluation of the ATCC 20-strain microbiome standard. (A) Presence-score ranking of the top candidate entries after evaluating the ATCC 20-strain reference set together with the UHGG representative genome background. Nineteen of the 20 reference taxa were significant at q ≤ 0.05. The remaining reference taxon, Propionibacterium acnes, was ranked at position 20 but did not pass the adjusted significance threshold. (B) Screening context showing the number of candidate entries, reference taxa, reference taxa called at q ≤ 0.05, and non-reference calls at q ≤ 0.05.


Supplementary Figure S5. Reference omission and additional genome calls. (A, B) Additional reference-genome calls at BH-adjusted q ≤ 0.01 after omission of each expected reference in SIHUMIx (A; eight scenarios) and pooled Mix24X near-genus (B; 24 scenarios). Bars distinguish calls already significant in the matching baseline (grey hatching) from newly significant calls after omission (orange). Numbers give total additional calls. All scenarios recovered the remaining expected genomes: 7/7 in SIHUMIx and 23/23 in Mix24X. Both panels use the same count scale and are ordered by total additional calls, with alphabetical tie-breaking. Reference annotations are retained; strain identifiers are omitted from display labels but retained in the source data. (C) Paired panel-unique peptide counts for the 73 SIHUMIx and 59 Mix24X newly significant call events. Each line links the same candidate genome before and after omission in one scenario; blue and orange points denote baseline and leave-out counts, respectively. The vertical coordinate is log10(count + 1), with ticks labelled in original count units. Small deterministic horizontal offsets separate overlapping events and have no biological meaning. Baseline unique counts were zero in 65/73 SIHUMIx and 52/59 Mix24X events (117/132 overall). Counts represent distinct I/L-normalized peptide sequences in the pooled inputs, not peptide-spectrum matches. Identified peptide inputs were held fixed; degeneracy, unique/shared classification, opportunity and scoring were recomputed. An event is a candidate genome in an omission scenario, not an independent biological observation or a distinct species. These comparisons do not establish peptide origins or biological presence. Raw component p-values and BH-adjusted q-values are retained in the paired source table and Supplementary Table S5; scenario summaries at q ≤ 0.05 are also supplied as source data.
Supplementary References

1.	Valdés-Mas, R., Leshem, A., Zheng, D., Cohen, Y., Kern, L., Zmora, N., He, Y., Katina, C., Eliyahu-Miller, S., and Yosef-Hevroni, T. (2025) Metagenome-informed metaproteomics of the human gut microbiome, host, and dietary exposome uncovers signatures of health and inflammatory bowel disease. Cell 188, 1062–1083. e1036
2.	Doellinger, J., Schneider, A., Hoeller, M., and Lasch, P. (2020) Sample preparation by easy extraction and digestion (SPEED)-a universal, rapid, and detergent-free protocol for proteomics based on acid extraction. Molecular & Cellular Proteomics 19, 209–222
